# Predicting the Pathway Involvement of Metabolites in Both Pathway Categories and Individual Pathways

**DOI:** 10.1101/2024.08.07.607025

**Authors:** Erik D. Huckvale, Hunter N.B. Moseley

## Abstract

Metabolism is the network of chemical reactions that sustain cellular life. Parts of this metabolic network are defined as metabolic pathways containing specific biochemical reactions. Products and reactants of these reactions are called metabolites, which are associated with certain human-defined metabolic pathways. Metabolic knowledgebases, such as the Kyoto Encyclopedia of Gene and Genomes (KEGG) contain metabolites, reactions, and pathway annotations; however, such resources are incomplete due to current limits of metabolic knowledge. To fill in missing metabolite pathway annotations, past machine learning models showed some success at predicting KEGG Level 2 pathway category involvement of metabolites based on their chemical structure. Here, we present the first machine learning model to predict metabolite association to more granular KEGG Level 3 metabolic pathways. We used a feature and dataset engineering approach to generate over one million metabolite-pathway entries in the dataset used to train a single binary classifier. This approach produced a mean Matthews correlation coefficient (MCC) of 0.806 ± 0.017 SD across 100 cross-validations iterations. The 172 Level 3 pathways were predicted with an overall MCC of 0.726. Moreover, metabolite association with the 12 Level 2 pathway categories were predicted with an overall MCC of 0.891, representing significant transfer learning from the Level 3 pathway entries. These are the best metabolite-pathway prediction results published so far in the field.

## 1. Introduction

Metabolism is the set of biochemical reactions occurring within organisms or individual cells that act to sustain life. These reactions are commonly represented as a metabolic network, i.e., a graph with metabolite as nodes and reaction as edges, with some edges having directionality. Parts of the metabolic network are defined as metabolic pathways based on many decades of study, research, and teaching of metabolism (1–3). From these pathway definitions, metabolites are associated with specific pathways. Metabolic knowledgebases such as the Kyoto Encyclopedia of Genes and Genomes (KEGG) (4–6), MetaCyc (7), and Reactome (8) contain mappings between metabolites and associated pathways, but these mappings are incomplete and are very slow, tedious, and costly to discover. For this reason, accurately predicting the pathway involvement of detected metabolites would be a valuable capability across biological and biomedical research fields.

Various attempts have been made to develop machine learning models to predict metabolite-pathway mappings given information about a metabolite’s chemical structure, especially pathway mappings for KEGG Level 2 (L2) ‘Metabolism’ pathway categories (9–12). In several published attempts, the dataset used was discovered to be invalid (13). Thus, Huckvale et al created a new benchmark dataset of metabolite features and associated pathway mappings as labels, all derived from KEGG (14). The features in this benchmark dataset were generated using an atom coloring technique developed by Jin and Moseley (15). This work involved training a binary classifier for each pathway, the input being a representation of a metabolite and the output indicating whether or not the given metabolite is associated with the pathway (14). Each model was only capable of predicting a single pathway. Through the use of a novel feature and pathway engineering approach, the later work of Huckvale and Moseley (16) created a single binary classifier capable of predicting an arbitrary metabolite-pathway mapping based on a pathway chemical structure representation. The novel feature and pathway engineering approach involves a cross join of metabolite features with pathway features, creating a dataset that is 12 times larger than the original benchmark dataset: 68,196 metabolite-pathway pair entries vs 5,683 metabolite entries (16).

Collections of pathways can be organized into pathway categories and these categories can be organized into broader categories in a hierarchical manner. In the case of KEGG, pathways are organized into a hierarchy where the first level of the hierarchy includes ‘Metabolism’, the top-level (Level 1 or L1) category containing metabolic pathways. The second level, i.e. L2, in the hierarchy includes 12 pathway categories sometimes referred to as modules. These L2 pathway categories were the focus of past works (9–12,14,16). However, there are 172 Level 3 (L3) pathways that are more granular. When combined with the broader 12 L2 pathway categories, a set of 184 metabolic “pathways” is created. In this situation, it is neither practical to train individual models for each pathway nor is it practical to train a multi-output model with 184 outputs; because, most of the resulting datasets would have too few positive entries for effective training. Having a significantly lower proportion of positive examples often results in excellent accuracy but poor recall, precision, and related metrics (17).

The cross join technique introduced by Huckvale and Moseley (16) resolves these issues, since it is a single model with a single output and capable of accepting an indefinite number of pathways into the training dataset. Specifically, the resulting dataset from the cross join increases the amount of positive examples to a level needed for effective training. Moreover, we show in this work that while the L3 pathways perform worse when trained by themselves, their performance increases when trained with the L2 pathway categories in the same dataset and with the same model. Likewise, the L2 pathways also increase in performance when trained alongside the L3 pathways. Combining both the broader pathway categories with the more granular individual pathways result in an improved overall model performance. This demonstrates that it is more than possible to train a model to reliably predict mappings to both L2 pathway categories and L3 individual pathways.

## 2. Materials and Methods

Our work began using the same methods as Huckvale et al (14) to construct features representing metabolites using the atom coloring technique introduced by Jin and Moseley (15). Likewise, the molfiles and pathway mappings were obtained from KEGG using the kegg_pull Python package (18). Then the same cross join technique, as introduced by Huckvale and Moseley (16), was used to construct the metabolite and pathway feature pairs. The metabolite-to-pathway mappings are defined by the KEGG pathway hierarchy available at: https://www.genome.jp/kegg-bin/show_brite?br08901.keg. We see that the L1 pathway categories in the hierarchy include ‘Metabolism’ and below that are the L2 pathway categories used in the prior works of Huckvale et al (14,16) and other publications involving metabolic pathway prediction with KEGG (9,10)(11,12). An additional hierarchy level below that, we see the L3 pathways are individual pathways as compared to categories of pathways in the hierarchy levels above. However, whether an individual pathway or a category of pathways, either can be defined as a collection of associated metabolites, enabling the construction of a cross joined dataset of combined metabolite and pathway features including both the L2 pathway categories and the L3 individual pathways. The L2 dataset was constructed by the prior work of Huckvale and Moseley (16). In this work, we constructed the L3 dataset containing the metabolic pathways in the 3^rd^ level of the KEGG hierarchy and the metabolites associated with them. Additionally, we constructed a 3^rd^ combined dataset containing both L2 and L3 pathways. Table 1 provides details of these three datasets, including the total number of entries (metabolite pathway pairs) after the cross join.

**Table 1.**
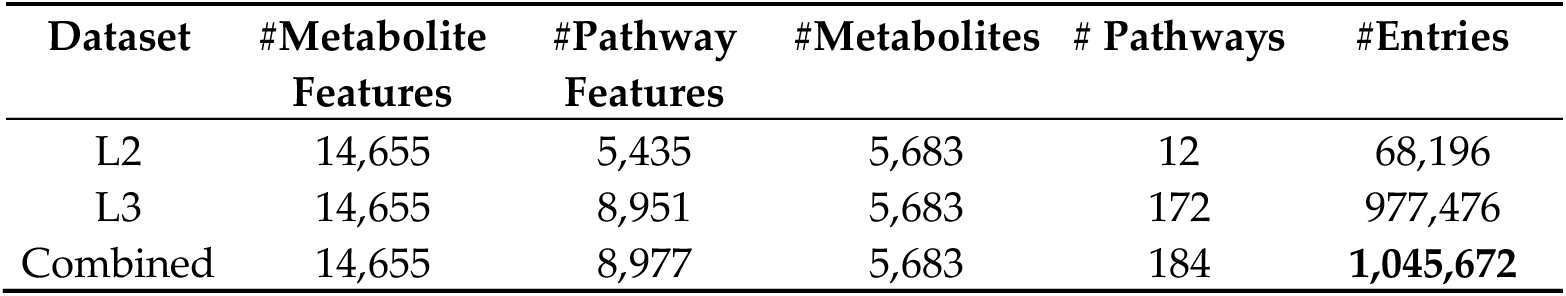
Description of cross joined metabolite-pathway paired datasets used for model training and testing.

Both metabolite and pathway features were normalized using the softmax by bond count technique performed in Huckvale and Moseley (16). While this is only an entrywise normalization, in this work, we additionally normalized feature-wise using standard min/max scaling (each feature value ranges between 0 and 1). To compare how the scaling impacts model performance, we created both a scaled and non-scaled version of each of the L2 only, L3 only, and combined datasets resulting in six total datasets. Note that the non-scaled L2 only dataset was part of the prior work of Huckvale and Moseley (16) and the five additional datasets were introduced in this work.

In the work of Huckvale and Moseley, the Multi-Layer Perceptron (MLP) model performed best and is also scalable, being able to perform just as well when training on the data in batches as opposed to current implementations of XGBoost (19), which are most optimal when trained on the entire dataset all at once (16). This is a limitation when the dataset does not fit entirely into memory, necessitating batched training. Considering the superior performance and ability to scale to indefinitely large datasets, we only trained and evaluated MLP models in this work. Models were trained to perform binary classification, i.e. given both metabolite features and pathway features, predict whether or not the given metabolite is associated with the given pathway. Hyperparameters were tuned for each of the six subsets, producing six corresponding models. Note that the model trained on the non-scaled L2 only dataset was part of the prior work of Huckvale and Moseley (16) and the five additional models were introduced in this work.

Models were evaluated using cross validation (CV) and stratified train test splits (20) over 100 CV iterations with 10% of the entries sampled in the test sets. Reducing the number of CV iterations from 1,000 to 100 was a pragmatic decision given the jump from training with 68,196 entries to 1,045,672 entries. Metrics used to evaluate the models included accuracy, precision, recall, F1 score, and Matthews correlation coefficient (MCC) (21). While these metrics were used to compare the test sets’ predictions and ground truth, we additionally counted the individual true positives, true negatives, false positives, and false negatives of each individual pathway. Meaning, we calculated the metrics across all entries in the test set of each CV iteration and we also split the test sets up by pathway category since each entry includes a pathway identity in the pair. We did not calculate these metrics for each pathway individually in each CV iteration because some of the pathways were too small i.e. did not have a sufficient number of metabolites associated with them (positive examples) in order to consistently calculate valid scores (zero true/false positives can cause a division by zero when calculating MCC). Counting the individual number of true positives, false negatives, true negatives, and false negatives, enabled us to sum these metrics across all 100 CV iterations and construct a single confusion matrix used to calculate the overall MCC per pathway. Since we were able to calculate metrics for the entire test set on each CV iteration, we were able to calculate means and standard deviations for the scores across all pathways. But individual pathway scores do not have standard deviations since only a single MCC score can be calculated from the confusion matrix summed across all the CV iterations.

The hardware used for this work included machines with up to 64 gigabytes (GB) of random-access memory (RAM) and central processing units (CPUs) ranging from 3.5 gigahertz (GHz) to 3.6 GHz, each with 6 hyperthreaded (HT) cores. The CPU chips included ‘Intel(R) Core(TM) i7-6850K CPU@3.60GHz’ and ‘Intel(R) Core(TM) i7-5930K CPU@3.50GHz’. The graphic processing units (GPUs) used had 16 GB of GPU RAM, with the name of the GPU card being ‘NVIDIA GeForce RTX 4080’.

All code for this work was written in major version 3 of the Python programming language (22). Data processing and storage was done using the Pandas (23), NumPy (24), and H5Py (25) packages. Models were constructed and trained using the PyTorch Lightning package (26) built upon the PyTorch package (27). Hyperparameters were tuned using the Optuna package (28). Metrics and the stratified train test splits were computed using the Sci-Kit Learn package (29). Results were stored in an SQL database (30) using the DuckDB package (31). Data visualizations were produced using the Tableau business intelligence software (32) as well as the Seaborn package (33) built on the MatPlotLib package (34). Statistical correlations were calculated using the SciPy package (35). Computational resource usage of the training and evaluation was collected using the gpu_tracker package (36). All code and data for reproducing these analyses can be accessed via the following Figshare item: https://doi.org/10.6084/m9.figshare.26505907.

## 3. Results

### 3.1. Overall Model Performance Across All Pathways

Figure 1 shows violin plots of the MCC scores across all 100 CV iterations for the L3 only and L2+L3 combined datasets, separated by scaled and non-scaled features. Each MCC score was computed from all predictions in the test set of each CV iteration. As we can see, the L2+L3 combined dataset has a higher median MCC and less variance than the L3 only dataset, whether the dataset was scaled or not. Scaling the features increased the median performance in both the L3 only and L2+L3 combined datasets, respectively.

**Figure 1.**
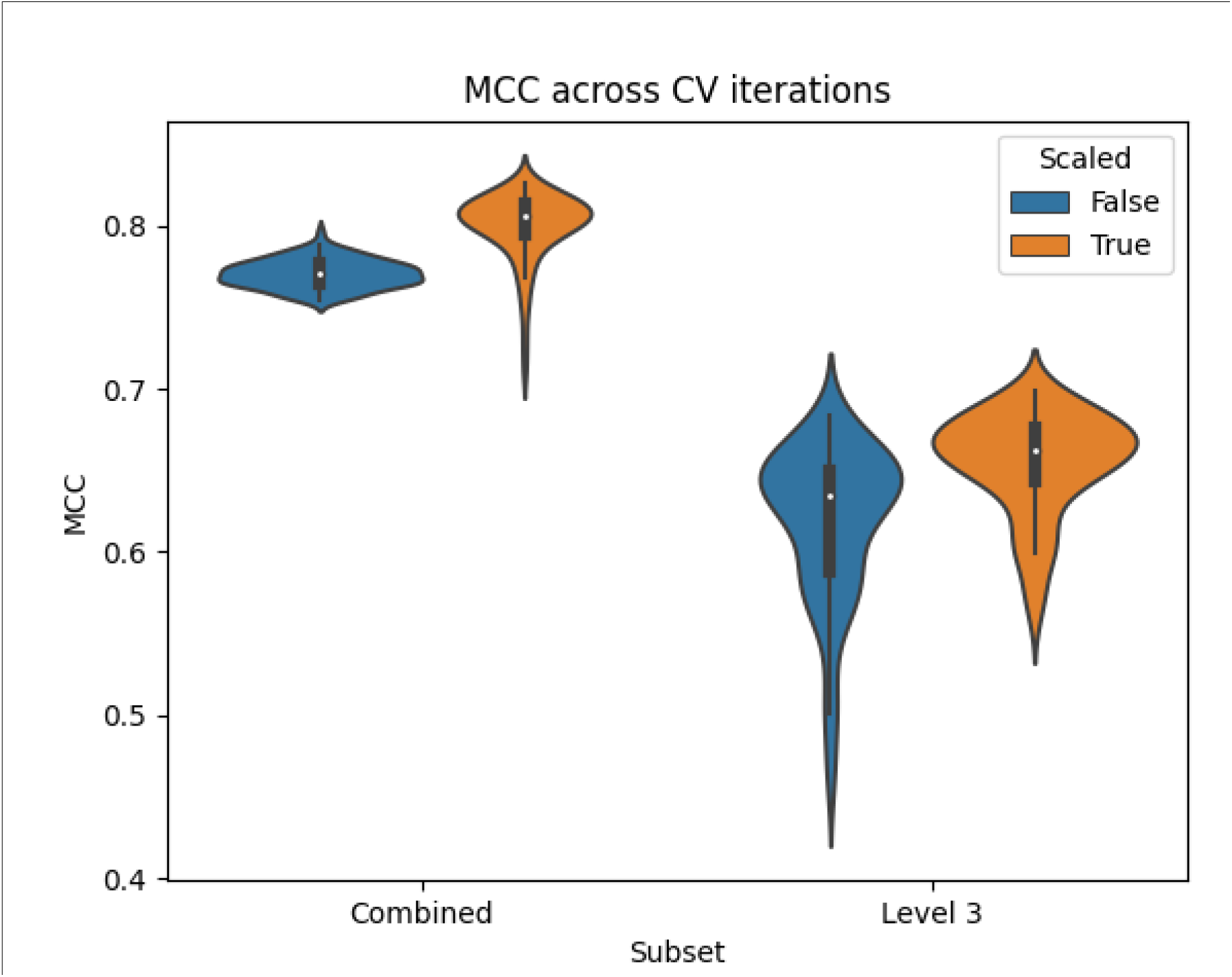
Violin plots of MCC scores across 100 CV iterations for models trained on the L3 only or L2+L3 combined datasets, with or without scaling.

Table 2 shows the mean and standard deviation MCC from the results in Figure 1 and additionally includes those for the L2 only dataset. Table S1 contains scores for the other 4 metrics i.e. accuracy, precision, recall, and F1 score. We see from Table 2 that while scaling the features improved the MCC of the L3 only and combined pathways, it decreased the MCC of the L2 only pathways. However, these MCCs are calculated from the entries available in the given dataset, with L2 entries missing from the L3 only dataset. So the comparisons of these MCCs and violin plots in Figure 1 are not exactly apples-to-apples and oranges-to-oranges comparisons.

**Table 2.** Mean model MCC performance metrics based on the given dataset.

| Dataset | Scaled | Mean MCC | Standard Deviation |
| --- | --- | --- | --- |
| Combined | True | **0.800** | 0.021 |
|  | False | 0.771 | 0.009 |
| L2 only | True | 0.728 | 0.029 |
|  | False | 0.784 | 0.013 |
| L3 only | True | 0.655 | 0.031 |
|  | False | 0.618 | 0.048 |

To address this shortcoming, Table 3 shows the MCC across only the L2 or L3 pathways within the L2+L3 combined dataset. While Table 2 shows the MCC of the L2 pathways after training only on the L2 dataset and the L3 pathways after training on only the L3 dataset, Table 3 shows the overall MCC specific to L2 and L3 pathways, separating them after training on the L2+L3 combined dataset. The MCCs in Table 3 were calculated by summing the true/false positives/negatives across all pathways of the given hierarchy level and across all CV iterations as described in the Methods, which prevents the calculation of a standard deviation. Table S2 contains these sums as well as the number of pathway / CV iteration pairs that were summed. Since there were 100 CV iterations and 12 L2 pathways, there were a total of 1,200 counts that contributed to the L2 sums. The L3 pathway sums likewise had 17,200 counts. The summed counts combined to construct confusion matrixes to calculate a single MCC for each combination of hierarchy level and whether or not the features were scaled. We see that for both hierarchy levels, the pathway categories predicted significantly better when trained on the combined dataset than when trained separately. While the non-scaled features performed better for the L2 pathways when training on the L2 pathways alone (Table 2), scaling resulted in greater performance for the L2 pathways when trained on the L2+L3 combined dataset (Table 3). Taking the best results between scaled vs unscaled datasets, the L2 only trained model had a mean MCC 0.784 vs an overall MCC of 0.891 for the L2+L3 trained model, representing a huge improvement!

**Table 3.** Overall L2+L3 combined model MCC performance metrics specific to pathway hierarchy level.

| Test set | Scaled | MCC |
| --- | --- | --- |
| L2 only | True | **0.891** |
|  | False | 0.850 |
| L3 only | True | 0.726 |
|  | False | 0.703 |

Table 4 shows the computational resource usage of the models trained on the L3 only and L2+L3 combined datasets, separated by whether the features were scaled. These are the computational resources used when evaluating the models over the 100 CV iterations. We see that scaling the features resulted in longer compute time but less GPU RAM and RAM. For the scaled datasets, the L3 only dataset took longer to train and evaluate despite there being less RAM utilization. This is likely due to the models requiring more epochs to converge. From Huckvale and Moseley (16), L2 only training and evaluation using an unscaled MLP model took roughly 90 minutes for 1000 CV iterations. In comparison, the training and evaluation on the L2+L3 combined unscaled dataset across 100 CV iterations took 131.1 hours. The large time increase in training and evaluation is to be expected since the L2 only dataset has 68,196 entries versus 1,045,672 entries in the L2+L3 combined dataset. Resource usage statistics were collected using the gpu_tracker package (36).

**Table 4.** Hardware utilization for model training and evaluation across 100 CV iterations.

| Dataset | Scaled | Resource | Unit | Amount |
| --- | --- | --- | --- | --- |
| Combined | True | Compute time | Hours | 131.8 |
|  |  | GPU RAM | Gigabytes | 4.3 |
|  |  | RAM | Gigabytes | 13.6 |
|  | False | Compute time | Hours | 131.1 |
|  |  | GPU RAM | Gigabytes | 4.6 |
|  |  | RAM | Gigabytes | 14.1 |
| L3 only | True | Compute time | Hours | 171.3 |
|  |  | GPU RAM | Gigabytes | 2.9 |
|  |  | RAM | Gigabytes | 12.6 |
|  | False | Compute time | Hours | 127.8 |
|  |  | GPU RAM | Gigabytes | 4.1 |
|  |  | RAM | Gigabytes | 16.6 |

### 3.2. Model Performance Per Pathway in the Combined Dataset

The following results are based solely on models trained on the scaled L2+L3 combined dataset. Table S3 contains the pathway information used to produce the following plots.

#### 3.2.1. Distribution of pathway statistics

There are two ways we measured the size of a pathway. The first is simply the number of metabolites associated with it. Each one of those metabolites have a certain number of non-hydrogen atoms as part of their molecular structure. So, the second way we measured a pathway size is the sum of all the non-hydrogen atoms across all the metabolites associated with the pathway. Figure 2 displays the distribution of the pathway sizes using both metrics.

**Figure 2.**
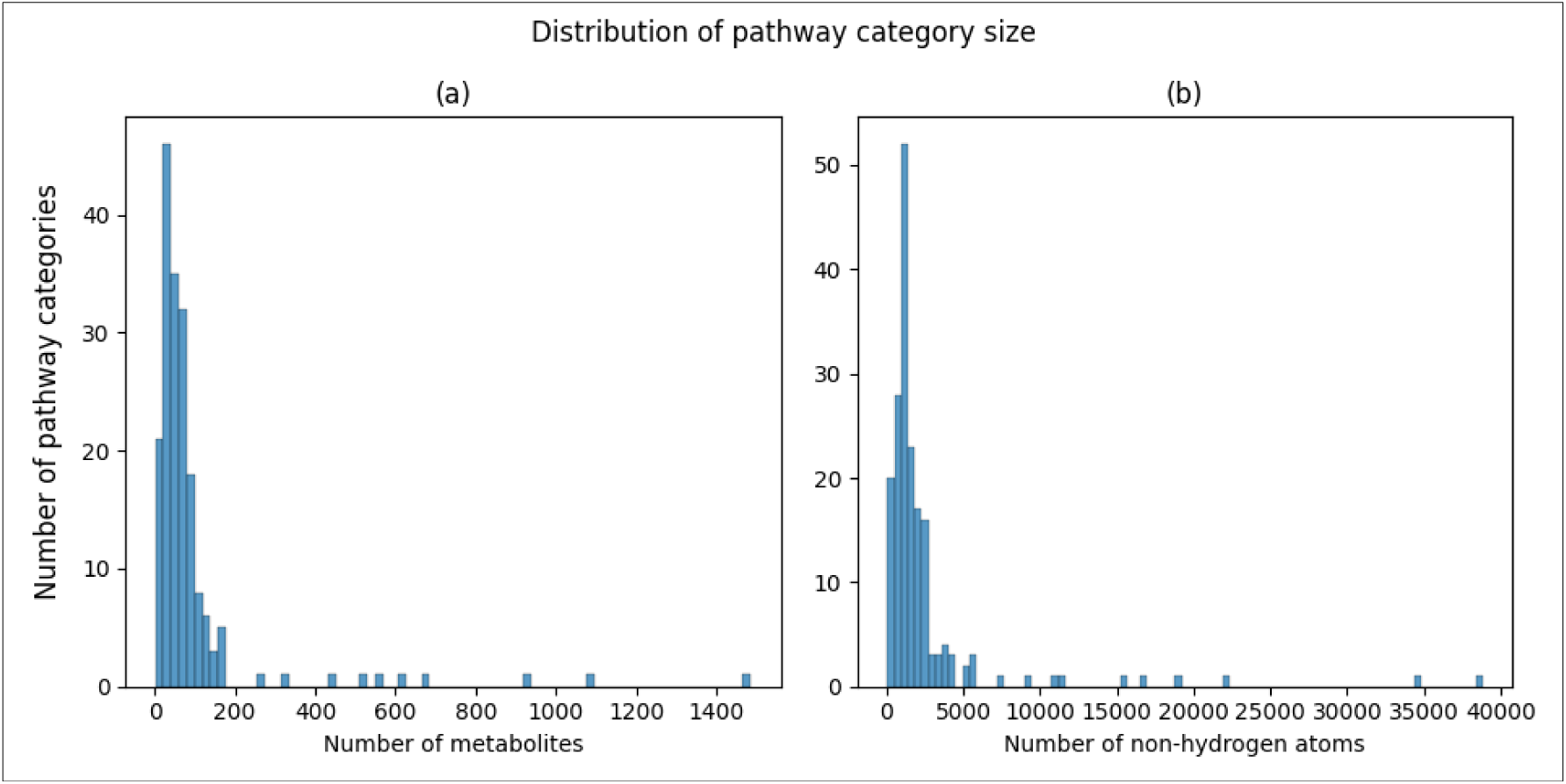
Histograms of pathway size. a) Histogram of number of metabolites associated with a pathway. b) Histogram of number of non-hydrogen atoms summed across metabolites associated with a pathway.

Figure 3 shows the distribution of overall pathway-specific MCC scores across 100 CV iterations. While 5 pathways scored an overall MCC of less than 0.4, the majority of pathways scored above 0.4, with a median of 0.722.

**Figure 3.**
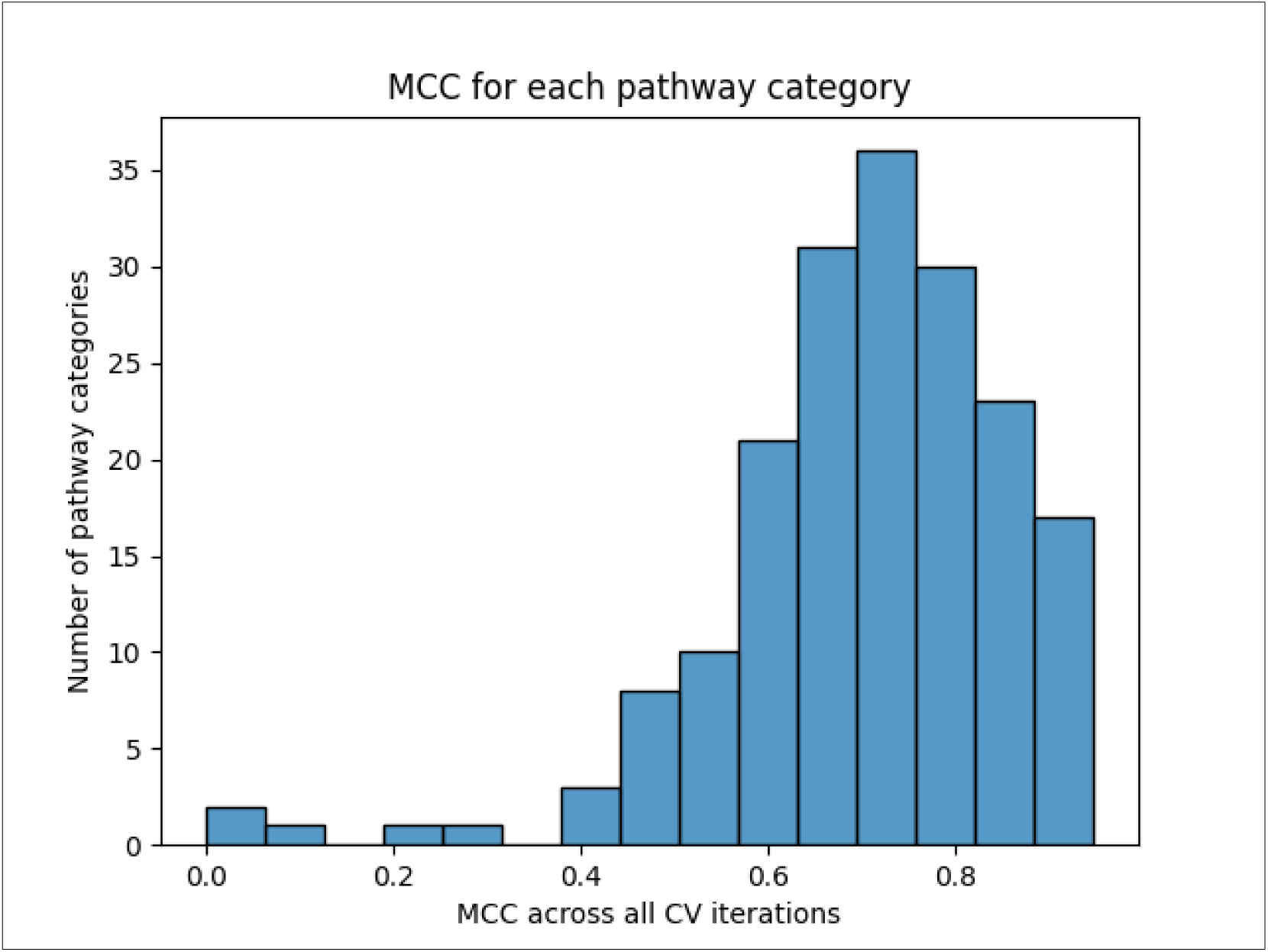
Histogram of pathway-specific overall MCC scores.

#### 3.2.2. Comparing pathway category size to MCC

Figure 4 shows scatterplots comparing the pathway size to its overall MCC score across 100 CV iterations. The X axes are shown both on the regular scale as well as on log10 scale. And both forms of measuring pathway size are shown. The log10 scale shows a clear nearly linear relationship between pathway size and MCC.

**Figure 4.**
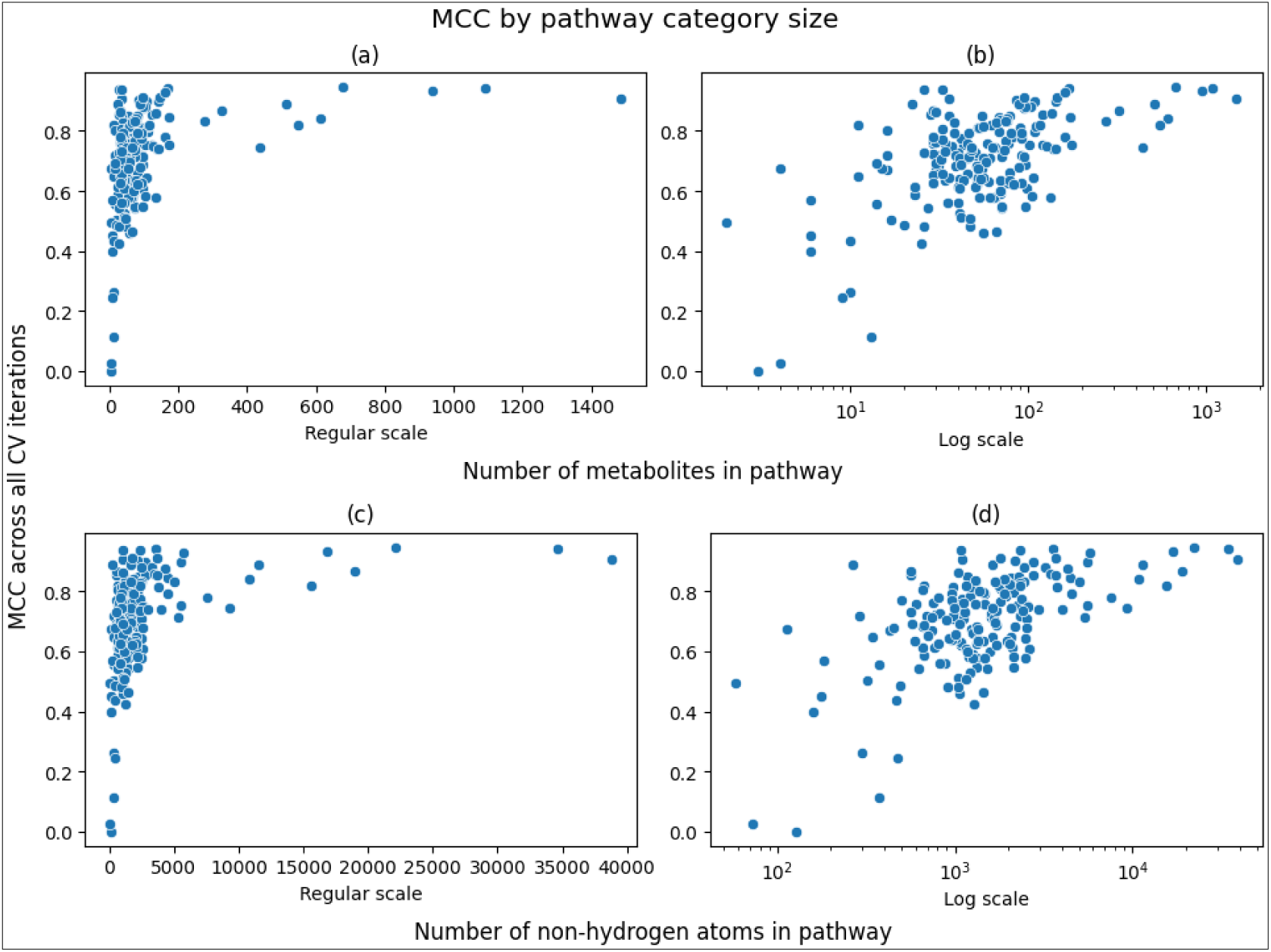
Scatterplots of overall MCC scores versus pathway size. a/b) Scatterplot of overall MCC score versus number of metabolites associated with a pathway. c/d) Scatterplot of overall MCC score versus number of non-hydrogen atoms summed across metabolites associated with a pathway.

Table 5 shows of the correlations between the size of the pathway to the overall MCC of the pathway, as plotted in Figure 4 above. While the correlation coefficients are moderate, the p-values are very low. We see that the correlation is higher and more statistically significant when measuring the pathway size as the sum of the number of non-hydrogen atoms across all compounds in the pathway as compared to the mere number of compounds in the pathway category. And the correlation is even stronger when calculating the Pearson correlation coefficient on the log10 scale compared to Spearman on the regular scale.

**Table 5.** Correlations between overall MCC and pathway size.

| Pathway size metric | Correlation Coefficient / Scale | p-Value |
| --- | --- | --- |
| # Compounds | 0.442 Spearman / regular | $3.463 \times 10^{-11}$ |
| # Non-hydrogen Atoms | 0.501 Spearman / regular | $4.234 \times 10^{-14}$ |
| # Compounds | 0.572 Pearson / log10 | $2.177 \times 10^{-17}$ |
| # Non-hydrogen Atoms | 0.584 Pearson / log10 | $3.160 \times 10^{-18}$ |

Since we can calculate the overall MCC of an individual pathway across the CV iterations, we can also calculate the overall MCC across subsets of pathways by summing their confusion matrixes. Figure 5 shows the increase of MCC across the remaining pathways after filtering pathways below a size threshold from the MCC calculation. For example, a threshold size of 0 is the overall MCC across all pathways and a threshold of 200 only calculates the MCC of pathways with at least 200 associated metabolites (Figure 5a). The same trend is shown when measuring pathway size by total number of non-hydrogen atoms with thresholds ranging up to 5,000 (Figure 5c). Filtering pathways from the calculation also results in less pathways to consider, and Figure 5b and 5d show the drop in remaining pathways as the threshold increases. We see an increase of MCC up to above 0.88 when only considering the 25 largest pathways. These trends are expected and simply highlight the higher information content of larger pathways with respect to model training.

**Figure 5.**
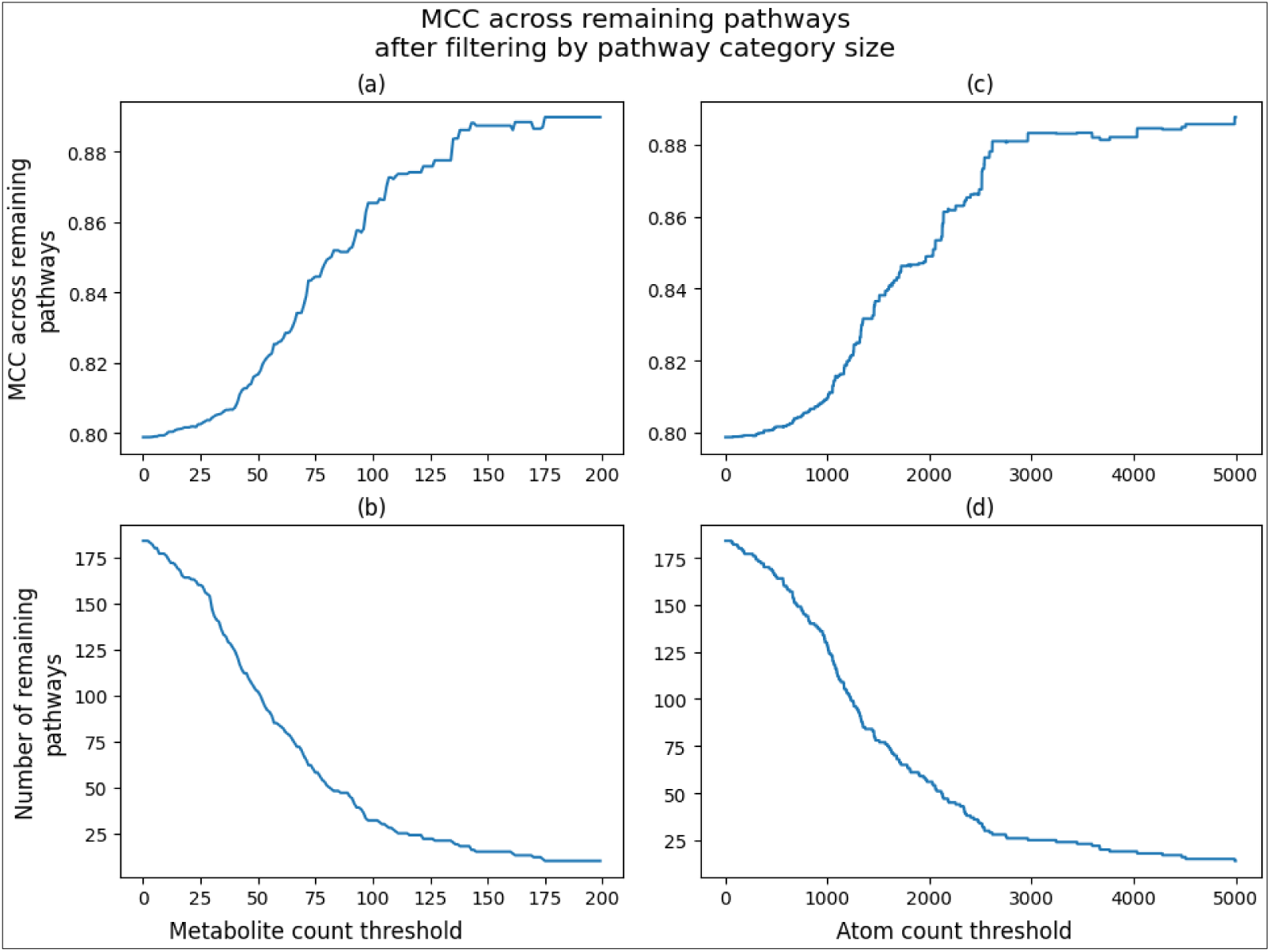
Line plots of overall MCC and pathway size based on pathway size filtering. a) Line plot of overall MCC based on metabolite count filtering. b) Line plot of number of pathways based on metabolite count filtering. c) Line plot of overall MCC based on non-hydrogen atom count filtering. d) Line plot of number of pathways based on non-hydrogen atom count filtering.

### 3.3. Impact of MCC when filtering pathways from the training set by pathway size thresholds

To determine how including smaller pathways (pathways with less associated metabolites) in the training set impacts the performance of the larger pathways, we created additional subsets by filtering pathways below a size threshold from the combined and scaled subset. With a threshold of 3, no metabolites were filtered since the smallest pathways had at least three metabolites associated with them. Using a threshold of 5 resulted in a subset containing 181 out of the 184 combined pathways and so on with thresholds of 7, 10, 12, 20, 50, and 100. This is distinct from the filtering in Figure 5 since those results were from filtering pathways from the MCC calculation, but the model was trained on all pathways. For the results in Figure 6, the model was trained on a subset of the pathways to determine how excluding pathways from the training set, not just the MCC calculation, impacts the performance of the 33 largest pathways. The largest threshold was 100, resulting in only 33 pathways remaining in its corresponding training set (Figure 6a). To make all of the filtering thresholds comparable, all MCCs reported are based on these 33 largest pathways. We see that pathways smaller than 12 hinder the performance of these 33 largest pathways with the lowest performance of 0.853 with a filter of 5 versus the highest performance of 0.872 with the 12-metabolite threshold. But filtering at 20 or above resulted in decreasing performance starting at 0.851.

**Figure 6.**
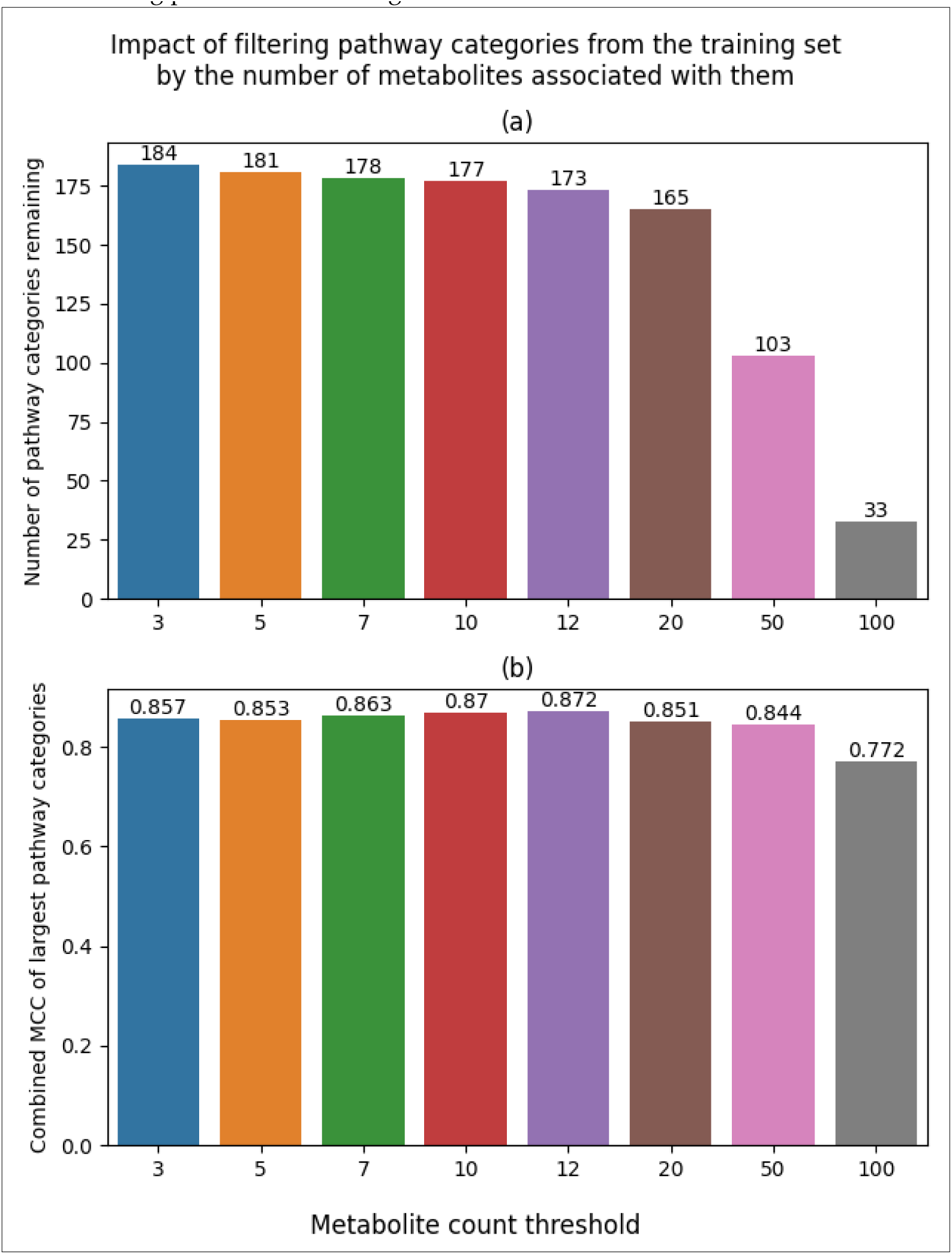
Bar plots of model performance based on pathway size filtering of the training set. a) Bar plot of the number of pathways included in training based on metabolite count filtering. b) Bar plot of overall MCC of the 33 largest pathways based on metabolite count filtering of the training set.

Since the 12-metabolite count threshold dataset performed the best, Figure 7 shows the distribution of MCC scores across CV iterations for the entirety of the test set containing 173 out of 184 original pathways. We see a 0.006 improvement (increase) in mean and 0.004 improvement (decrease) in standard deviation compared to the scaled L2+L3 combined dataset that contained all 184 pathways (Table 2).

**Figure 7.**
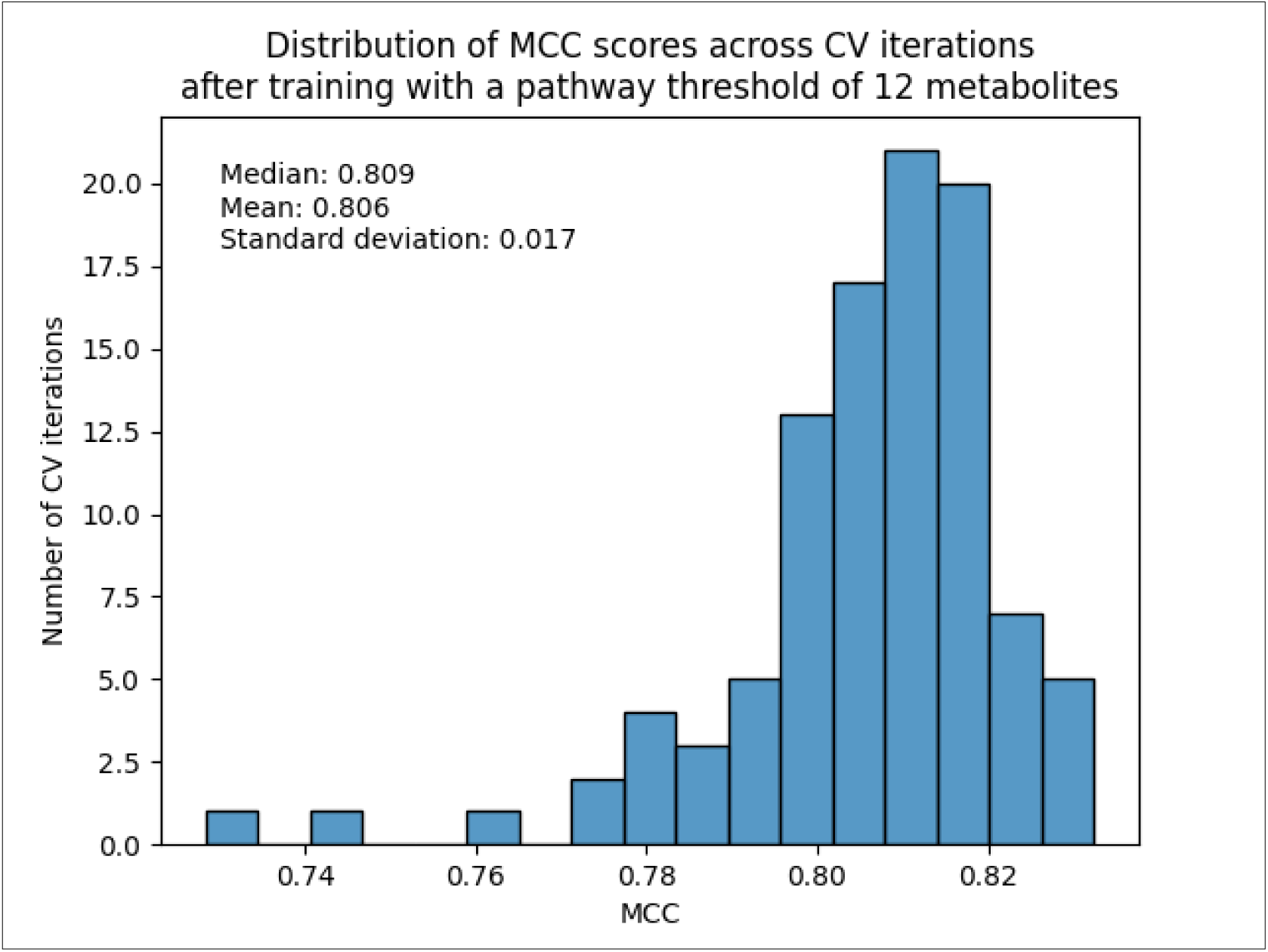
Histogram of MCC score across 100 CV iterations trained on the 12-metabolite count threshold dataset.

## 4. Discussion

In this work, we demonstrate that it’s possible to develop a single binary classifier that can predict not just on KEGG L2 metabolic pathway categories, but more granular L3 pathways, with MCCs mostly above 0.4 and reaching above 0.8 for the largest pathways. Like the work in Huckvale and Moseley (16), this single binary classifier approach uses a cross join of metabolite and pathway features to create a much larger dataset for training and testing, while also allowing prediction on arbitrary metabolites and pathways. This approach enabled the prediction of not just the 12 L2 pathways but the 184 L2 + L3 combined pathways by eliminating the complexity and limitations of training and deploying a binary classifier for every pathway category or a multi-output model with an unrealistically high number of outputs. In particular, a multi-output model would need to be retrained with each new pathway added.

Models trained only on L3 pathways did not perform as well (mean MCC of 0.655 ± 0.031 SD) as models trained only on L2 pathways (mean MCC of 0.784 ± 0.013 SD). Models trained on the L2+L3 combined dataset outperformed all previous models with a mean MCC of 0.800 ± 0.021 SD. Moreover, when just looking at performance of L2 pathways, the L2+L3 combined model had an overall MCC of 0.891! These results clearly demonstrate that adding more pathways increases the overall performance of the model, especially for large pathways. These results are a clear example of transfer learning, where training on additional pathways improve the performance for other pathways. However, we observed a clear trend where smaller pathways do perform worse than larger pathways. But there appears to be a sweet spot around pathways with 12 metabolites and larger that add more information than noise during model training. A training set trimmed to pathways with 12 or more metabolites maximized the overall MCC of 0.872 for the 33 largest pathways as compared to trainsets with more or fewer pathways. This final 12-metabolite threshold dataset produced a mean MCC of 0.806 ± 0.017 SD. Therefore, we recommend that future datasets filter pathways that are too small from the training set in order to improve the performance of the remaining pathways. However, the model is still capable of predicting on these smaller pathways if desired. We also observe an increase in performance when normalizing the data feature-wise in addition to entry-wise. Thus, we recommend min-max scaling for future datasets in addition to filtering by pathway size.

## 5. Conclusions

Despite their increased difficulty to predict, this work demonstrates that metabolite association with individual L3 pathways can be predicted with an overall MCC of 0.726. Moreover, metabolite association with L2 pathways can be predicted with an overall MCC of 0.891 using the L2+L3 combined dataset, representing transfer learning with increased dataset size. In general, larger pathways perform better than smaller pathways and the smallest pathways can even decrease the performance of larger pathways. The 12-metabolite threshold dataset produced the highest mean MCC of 0.806 ± 0.017 SD. Thus, filtering these smallest pathways can maximize model performance but filtering too much can decrease the performance due to removing valuable information for training.

## Supplementary Materials

Table S1: Model performance metrics for each dataset; Table S2: Sum of True/False Positives/Negative metrics across CV iterations by pathway hierarchy level (L2 or L3); Table S3: Characteristics and performance metrics for each L2 and L3 pathway.

## Author Contributions

Conceptualization, E.D.H and H.N.B.M.; methodology, E.D.H.; software, E.D.H.; validation, E.D.H.; formal analysis, E.D.H.; investigation, E.D.H.; resources, E.D.H.; data curation, E.D.H.; writing—original draft preparation, E.D.H.; writing—review and editing, E.D.H. and H.N.B.M.; visualization, E.D.H.; supervision, H.N.B.M.; project administration, H.N.B.M.; funding acquisition, H.N.B.M. All authors have read and agreed to the published version of the manuscript.

## Institutional Review Board Statement

Not applicable

## Informed Consent Statement

Not applicable

## Data Availability Statement

All code and data for reproducing these analyses can be accessed via the following Figshare item: https://doi.org/10.6084/m9.figshare.26505907.

## Acknowledgments

We thank the University of Kentucky Center for Computational Sciences and Information Technology Services Research Computing for their support and use of the Lipscomb Compute Cluster and associated research computing resources.

---

**Table S1.** Model performance metrics for each dataset.

| Subset | Scaled | Metric | Mean Score | Standard deviation |
| --- | --- | --- | --- | --- |
| Combined | True | Accuracy | 0.9930 | 0.0011 |
|  |  | F1 score | 0.8013 | 0.0223 |
|  |  | MCC | 0.7997 | 0.0209 |
|  |  | Precision | 0.7551 | 0.0396 |
|  |  | Recall | 0.8551 | 0.0129 |
|  | False | Accuracy | 0.9926 | 0.0003 |
|  |  | F1 score | 0.7743 | 0.0085 |
|  |  | MCC | 0.7706 | 0.0086 |
|  |  | Precision | 0.7735 | 0.0176 |
|  |  | Recall | 0.7754 | 0.0130 |
| Level 2 | True | Accuracy | 0.9473 | 0.0082 |
|  |  | F1 score | 0.7556 | 0.0262 |
|  |  | MCC | 0.7275 | 0.0294 |
|  |  | Precision | 0.7528 | 0.0575 |
|  |  | Recall | 0.7629 | 0.0340 |
|  | * False | Accuracy | 0.9592 | 0.0027 |
|  |  | F1 score | 0.8069 | 0.0114 |
|  |  | MCC | 0.7844 | 0.0129 |
|  |  | Precision | 0.8128 | 0.0224 |
|  |  | Recall | 0.8019 | 0.0193 |
| Level 3 | True | Accuracy | 0.9930 | 0.0015 |
|  |  | F1 score | 0.6529 | 0.0359 |
|  |  | MCC | 0.6547 | 0.0314 |
|  |  | Precision | 0.6830 | 0.0883 |
|  |  | Recall | 0.6440 | 0.0821 |
|  | False | Accuracy | 0.9902 | 0.0041 |
|  |  | F1 score | 0.6101 | 0.0628 |
|  |  | MCC | 0.6182 | 0.0481 |
|  |  | Precision | 0.5593 | 0.1238 |
|  |  | Recall | 0.7129 | 0.0787 |
| * The results highlighted in yellow were provided by the work of Huckvale and Moseley in the supplemental materials of the manuscript of DOI: https://doi.org/10.3390/metabo14050266 |  |  |  |  |

**Table S2.** Sum of True/False Positives/Negative metrics across CV iterations by pathway hierarchy level (L2 or L3). Results are from training on the combined and scaled dataset.

| Subset | Pathway Hierarchy Level | Scaled | Metric | Sum | Count (number of pathway categories times 100 CV iterations) |
| --- | --- | --- | --- | --- | --- |
| Combined | 2 | TRUE | True positives | 69,342 | 1,200 |
|  |  |  | True negatives | 597,569 | 1,200 |
|  |  |  | False positives | 12,142 | 1,200 |
|  |  |  | False negatives | 2,963 | 1,200 |
|  |  | FALSE | True positives | 63,596 | 1,200 |
|  |  |  | True negatives | 599,093 | 1,200 |
|  |  |  | False positives | 10,963 | 1,200 |
|  |  |  | False negatives | 8,677 | 1,200 |
|  | 3 | TRUE | True positives | 77,484 | 17,200 |
|  |  |  | True negatives | 9,639,265 | 17,200 |
|  |  |  | False positives | 36,124 | 17,200 |
|  |  |  | False negatives | 21,911 | 17,200 |
|  |  | FALSE | True positives | 69,546 | 17,200 |
|  |  |  | True negatives | 9,646,907 | 17,200 |
|  |  |  | False positives | 28,137 | 17,200 |
|  |  |  | False negatives | 29,881 | 17,200 |

**Table S3.** Characteristics and performance metrics for each L2 and L3 pathway. Results derive from training on the combined and scaled dataset.

| Pathway ID | Number of metabolites | Number of non-hydrogen atoms | MCC score | True positives | True negatives | False positives | False negatives |
| --- | --- | --- | --- | --- | --- | --- | --- |
| Amino acid metabolism | 611 | 10774 | 0.8409598893071340 | 5792.0 | 49225.0 | 1555.0 | 391.0 |
| Biosynthesis of other secondary metabolites | 1486 | 38836 | 0.9081988788786460 | 14491.0 | 40591.0 | 1735.0 | 382.0 |
| Carbohydrate metabolism | 514 | 11485 | 0.8902161978132910 | 4837.0 | 50831.0 | 898.0 | 195.0 |
| Chemical structure transformation maps | 437 | 9298 | 0.7442351678288250 | 3885.0 | 50243.0 | 1999.0 | 483.0 |
| Energy metabolism | 173 | 4470 | 0.8441864642903980 | 1539.0 | 54626.0 | 398.0 | 157.0 |
| Glycan biosynthesis and metabolism | 324 | 18936 | 0.8687650948335800 | 3133.0 | 52902.0 | 730.0 | 172.0 |
| Lipid metabolism | 677 | 22159 | 0.9480287144475790 | 6788.0 | 49618.0 | 535.0 | 119.0 |
| Metabolism of cofactors and vitamins | 549 | 15569 | 0.8192698358391200 | 5099.0 | 49542.0 | 1800.0 | 269.0 |
| Metabolism of other amino acids | 274 | 4989 | 0.8341470754511110 | 2554.0 | 53293.0 | 760.0 | 217.0 |
| Metabolism of terpenoids and polyketides | 1093 | 34676 | 0.9402708096069460 | 10590.0 | 44929.0 | 756.0 | 317.0 |
| Nucleotide metabolism | 169 | 3592 | 0.9420564036585730 | 1650.0 | 55260.0 | 166.0 | 33.0 |
| Xenobiotics biodegradation and metabolism | 939 | 16780 | 0.9349955392946330 | 8984.0 | 46509.0 | 810.0 | 228.0 |
| 00010 Glycolysis / Gluconeogenesis | 43 | 997 | 0.7521514556232700 | 349.0 | 55995.0 | 129.0 | 99.0 |
| 00020 Citrate cycle (TCA cycle) | 42 | 1089 | 0.7736618071841690 | 346.0 | 56150.0 | 142.0 | 62.0 |
| 00030 Pentose phosphate pathway | 65 | 1189 | 0.5801059320016230 | 424.0 | 55899.0 | 425.0 | 194.0 |
| 00040 Pentose and glucuronate interconversions | 69 | 1256 | 0.5896851476187490 | 457.0 | 55574.0 | 410.0 | 223.0 |
| 00051 Fructose and mannose metabolism | 71 | 1506 | 0.5425171605375680 | 424.0 | 55891.0 | 370.0 | 328.0 |
| 00052 Galactose metabolism | 64 | 1330 | 0.578369297572232 | 410.0 | 55245.0 | 381.0 | 214.0 |
| 00053 Ascorbate and aldarate metabolism | 71 | 1325 | 0.5480372306228640 | 465.0 | 56062.0 | 517.0 | 251.0 |
| 00061 Fatty acid biosynthesis | 23 | 670 | 0.5857307522988000 | 202.0 | 56524.0 | 270.0 | 48.0 |
| 00062 Fatty acid elongation | 39 | 2179 | 0.6824945492484340 | 347.0 | 55567.0 | 339.0 | 27.0 |
| 00071 Fatty acid degradation | 51 | 2484 | 0.6439211962872030 | 437.0 | 56211.0 | 423.0 | 92.0 |
| 00073 Cutin, suberine and wax biosynthesis | 23 | 656 | 0.6142145231530840 | 185.0 | 56583.0 | 223.0 | 36.0 |
| 00100 Steroid biosynthesis | 68 | 2069 | 0.8438381240718750 | 616.0 | 55703.0 | 131.0 | 94.0 |
| 00120 Primary bile acid biosynthesis | 55 | 2299 | 0.8489302256003490 | 491.0 | 56404.0 | 124.0 | 51.0 |
| 00121 Secondary bile acid biosynthesis | 36 | 1184 | 0.8496184733002620 | 345.0 | 56321.0 | 80.0 | 42.0 |
| 00130 Ubiquinone and other terpenoid-quinone biosynthesis | 91 | 2339 | 0.7202567520407570 | 711.0 | 55463.0 | 313.0 | 228.0 |
| 00140 Steroid hormone biosynthesis | 108 | 2745 | 0.8985746926606160 | 984.0 | 55486.0 | 163.0 | 57.0 |
| 00190 Oxidative phosphorylation | 14 | 372 | 0.5561799510943540 | 81.0 | 56711.0 | 88.0 | 44.0 |
| 00195 Photosynthesis | 6 | 184 | 0.5687909308210510 | 43.0 | 56722.0 | 49.0 | 19.0 |
| 00220 Arginine biosynthesis | 29 | 605 | 0.7568809031535640 | 214.0 | 56472.0 | 49.0 | 89.0 |
| 00230 Purine metabolism | 121 | 2755 | 0.8532967940978590 | 1212.0 | 55166.0 | 366.0 | 56.0 |
| 00232 Caffeine metabolism | 28 | 563 | 0.8541798746855700 | 242.0 | 56951.0 | 26.0 | 57.0 |
| 00240 Pyrimidine metabolism | 90 | 1787 | 0.8113729465430260 | 837.0 | 55378.0 | 293.0 | 97.0 |
| 00250 Alanine, aspartate and glutamate metabolism | 51 | 1075 | 0.6216987663625870 | 310.0 | 56068.0 | 188.0 | 184.0 |
| 00253 Tetracycline biosynthesis | 46 | 1468 | 0.7919846416467210 | 412.0 | 56190.0 | 192.0 | 34.0 |
| 00254 Aflatoxin biosynthesis | 36 | 1093 | 0.9055626098952720 | 337.0 | 56389.0 | 44.0 | 26.0 |
| 00260 Glycine, serine and threonine metabolism | 60 | 1074 | 0.632184821848565 | 397.0 | 56010.0 | 229.0 | 225.0 |
| 00261 Monobactam biosynthesis | 49 | 1098 | 0.7165231174376000 | 353.0 | 56369.0 | 142.0 | 134.0 |
| 00270 Cysteine and methionine metabolism | 96 | 1699 | 0.6882694148003660 | 721.0 | 55366.0 | 376.0 | 263.0 |
| 00280 Valine, leucine and isoleucine degradation | 52 | 1633 | 0.6458800086230890 | 393.0 | 56501.0 | 311.0 | 127.0 |
| 00290 Valine, leucine and isoleucine biosynthesis | 29 | 503 | 0.7687005955828100 | 241.0 | 56030.0 | 70.0 | 74.0 |
| 00300 Lysine biosynthesis | 55 | 1231 | 0.675042799256142 | 391.0 | 55833.0 | 222.0 | 151.0 |
| 00310 Lysine degradation | 70 | 1565 | 0.6013072452787590 | 428.0 | 55910.0 | 284.0 | 272.0 |
| 00311 Penicillin and cephalosporin biosynthesis | 31 | 692 | 0.7169593911073640 | 222.0 | 56107.0 | 83.0 | 91.0 |
| 00330 Arginine and proline metabolism | 77 | 1316 | 0.624971876536468 | 555.0 | 55507.0 | 468.0 | 202.0 |
| 00331 Clavulanic acid biosynthesis | 17 | 319 | 0.5018821415057870 | 90.0 | 56570.0 | 69.0 | 111.0 |
| 00332 Carbapenem biosynthesis | 45 | 1047 | 0.7562328936259140 | 348.0 | 56338.0 | 107.0 | 115.0 |
| 00333 Prodigiosin biosynthesis | 29 | 784 | 0.7242515796302220 | 227.0 | 56171.0 | 98.0 | 74.0 |
| 00340 Histidine metabolism | 61 | 1115 | 0.7109610395634370 | 449.0 | 55834.0 | 212.0 | 149.0 |
| 00350 Tyrosine metabolism | 95 | 1600 | 0.7132958932262360 | 720.0 | 55563.0 | 365.0 | 206.0 |
| 00360 Phenylalanine metabolism | 61 | 1463 | 0.5784794345723250 | 353.0 | 55826.0 | 288.0 | 219.0 |
| 00361 Chlorocyclohexane and chlorobenzene degradation | 91 | 1206 | 0.784074944506343 | 769.0 | 55253.0 | 249.0 | 167.0 |
| 00362 Benzoate degradation | 97 | 2614 | 0.6105447143364770 | 648.0 | 55191.0 | 490.0 | 319.0 |
| 00363 Bisphenol degradation | 29 | 568 | 0.869213773716127 | 240.0 | 56730.0 | 43.0 | 29.0 |
| 00364 Fluorobenzoate degradation | 40 | 817 | 0.7277840822453640 | 271.0 | 56572.0 | 81.0 | 121.0 |
| 00365 Furfural degradation | 16 | 428 | 0.6696526147960390 | 113.0 | 56955.0 | 55.0 | 56.0 |
| 00380 Tryptophan metabolism | 109 | 2126 | 0.7957896057720780 | 941.0 | 55135.0 | 262.0 | 209.0 |
| 00400 Phenylalanine, tyrosine and tryptophan biosynthesis | 56 | 1015 | 0.6432877086056760 | 373.0 | 56476.0 | 183.0 | 225.0 |
| 00401 Novobiocin biosynthesis | 37 | 1012 | 0.7470854287503790 | 298.0 | 56205.0 | 107.0 | 93.0 |
| 00402 Benzoxazinoid biosynthesis | 16 | 291 | 0.7176017065480430 | 102.0 | 56445.0 | 40.0 | 40.0 |
| 00403 Indole diterpene alkaloid biosynthesis | 38 | 1301 | 0.8302009318289570 | 325.0 | 56047.0 | 96.0 | 38.0 |
| 00404 Staurosporine biosynthesis | 64 | 1779 | 0.8191649844663780 | 539.0 | 56090.0 | 133.0 | 102.0 |
| 00405 Phenazine biosynthesis | 41 | 952 | 0.6381257912091950 | 280.0 | 56366.0 | 167.0 | 147.0 |
| 00410 beta-Alanine metabolism | 45 | 973 | 0.7087123261934690 | 340.0 | 56524.0 | 158.0 | 119.0 |
| 00430 Taurine and hypotaurine metabolism | 30 | 756 | 0.7607651217055730 | 237.0 | 56470.0 | 72.0 | 76.0 |
| 00440 Phosphonate and phosphinate metabolism | 68 | 1453 | 0.7966198293069850 | 572.0 | 56491.0 | 187.0 | 103.0 |
| 00450 Selenocompound metabolism | 31 | 655 | 0.6830059208336810 | 235.0 | 56240.0 | 151.0 | 70.0 |
| 00460 Cyanoamino acid metabolism | 50 | 763 | 0.6116918933903410 | 325.0 | 56103.0 | 233.0 | 175.0 |
| 00470 D-Amino acid metabolism | 79 | 1252 | 0.6598403086147240 | 563.0 | 55805.0 | 320.0 | 249.0 |
| 00480 Glutathione metabolism | 53 | 1258 | 0.5799322025862940 | 308.0 | 56044.0 | 260.0 | 182.0 |
| 00500 Starch and sucrose metabolism | 54 | 1346 | 0.6392912010311820 | 361.0 | 56104.0 | 249.0 | 156.0 |
| 00510 N-Glycan biosynthesis | 26 | 1129 | 0.7291468418447760 | 200.0 | 56482.0 | 94.0 | 55.0 |
| 00512 Mucin type O-glycan biosynthesis | 3 | 127 | -<br>0.00027401531865905300 | 0.0 | 56871.0 | 9.0 | 27.0 |
| 00513 Various types of N-glycan biosynthesis | 4 | 73 | 0.027205888688917800 | 1.0 | 56796.0 | 37.0 | 33.0 |
| 00514 Other types of O-glycan biosynthesis | 6 | 175 | 0.4525634034550170 | 24.0 | 56442.0 | 22.0 | 37.0 |
| 00515 Mannose type O-glycan biosynthesis | 6 | 158 | 0.39863663629348100 | 25.0 | 56606.0 | 45.0 | 31.0 |
| 00520 Amino sugar and nucleotide sugar metabolism | 137 | 4033 | 0.7401919779086580 | 1122.0 | 55133.0 | 547.0 | 230.0 |
| 00521 Streptomycin biosynthesis | 39 | 969 | 0.7195865469126920 | 294.0 | 56349.0 | 121.0 | 106.0 |
| 00522 Biosynthesis of 12-, 14- and 16-membered macrolides | 95 | 4281 | 0.8777338297470100 | 832.0 | 56092.0 | 137.0 | 91.0 |
| 00523 Polyketide sugar unit biosynthesis | 73 | 2432 | 0.8319233194762350 | 672.0 | 55971.0 | 223.0 | 53.0 |
| 00524 Neomycin, kanamycin and gentamicin biosynthesis | 103 | 3240 | 0.8811058059278500 | 892.0 | 55873.0 | 128.0 | 108.0 |
| 00525 Acarbose and validamycin biosynthesis | 52 | 1224 | 0.8125023355285470 | 446.0 | 55946.0 | 132.0 | 73.0 |
| 00531 Glycosaminoglycan degradation | 13 | 375 | 0.1117813630720040 | 13.0 | 56619.0 | 78.0 | 131.0 |
| 00532 Glycosaminoglycan biosynthesis - chondroitin sulfate / dermatan sulfate | 4 | 114 | 0.6764558122534930 | 25.0 | 56932.0 | 10.0 | 14.0 |
| 00534 Glycosaminoglycan biosynthesis - heparan sulfate / heparin | 2 | 58 | 0.49601170570954100 | 8.0 | 56530.0 | 18.0 | 2.0 |
| 00540 Lipopolysaccharide biosynthesis | 71 | 5332 | 0.7119255155083730 | 579.0 | 55988.0 | 300.0 | 165.0 |
| 00541 O-Antigen nucleotide sugar biosynthesis | 110 | 3764 | 0.8140831474534970 | 994.0 | 55373.0 | 376.0 | 84.0 |
| 00550 Peptidoglycan biosynthesis | 41 | 2517 | 0.7096009771486460 | 337.0 | 56391.0 | 199.0 | 81.0 |
| 00552 Teichoic acid biosynthesis | 40 | 2113 | 0.6129455314284880 | 292.0 | 56336.0 | 271.0 | 107.0 |
| 00561 Glycerolipid metabolism | 50 | 1585 | 0.7288252640151510 | 411.0 | 55887.0 | 202.0 | 104.0 |
| 00562 Inositol phosphate metabolism | 59 | 1456 | 0.7590277931409680 | 434.0 | 56088.0 | 130.0 | 142.0 |
| 00563 Glycosylphosphatidylinositol (GPI)-anchor biosynthesis | 10 | 298 | 0.2623363639353410 | 36.0 | 56544.0 | 148.0 | 65.0 |
| 00564 Glycerophospholipid metabolism | 76 | 1973 | 0.8329444010426630 | 686.0 | 55779.0 | 191.0 | 83.0 |
| 00565 Ether lipid metabolism | 45 | 1002 | 0.6583909820237230 | 317.0 | 56322.0 | 245.0 | 92.0 |
| 00571 Lipoarabinomannan (LAM) biosynthesis | 10 | 466 | 0.4353021170675020 | 55.0 | 56497.0 | 88.0 | 56.0 |
| 00572 Arabinogalactan biosynthesis - Mycobacterium | 9 | 474 | 0.2429860220833520 | 28.0 | 56063.0 | 113.0 | 65.0 |
| 00590 Arachidonic acid metabolism | 85 | 2186 | 0.9020556642167380 | 803.0 | 55749.0 | 144.0 | 31.0 |
| 00591 Linoleic acid metabolism | 30 | 744 | 0.6694539415743330 | 251.0 | 56492.0 | 171.0 | 80.0 |
| 00592 alpha-Linolenic acid metabolism | 62 | 2027 | 0.8213088985460710 | 529.0 | 55909.0 | 150.0 | 79.0 |
| 00600 Sphingolipid metabolism | 56 | 1650 | 0.7081600596698010 | 490.0 | 55922.0 | 329.0 | 89.0 |
| 00601 Glycosphingolipid biosynthesis - lacto and neolacto series | 26 | 2339 | 0.9355472092944740 | 255.0 | 56329.0 | 21.0 | 14.0 |
| 00603 Glycosphingolipid biosynthesis - globo and isoglobo series | 11 | 673 | 0.8199876238303620 | 93.0 | 56348.0 | 26.0 | 15.0 |
| 00604 Glycosphingolipid biosynthesis - ganglio series | 16 | 1368 | 0.8008036639157520 | 123.0 | 56686.0 | 29.0 | 32.0 |
| 00620 Pyruvate metabolism | 51 | 1324 | 0.6833139881448820 | 377.0 | 55811.0 | 189.0 | 156.0 |
| 00621 Dioxin degradation | 69 | 1294 | 0.8371759600896710 | 584.0 | 56071.0 | 102.0 | 122.0 |
| 00622 Xylene degradation | 46 | 740 | 0.7119955738202290 | 335.0 | 56375.0 | 127.0 | 141.0 |
| 00623 Toluene degradation | 56 | 1059 | 0.4611271845303230 | 272.0 | 55978.0 | 276.0 | 348.0 |
| 00624 Polycyclic aromatic hydrocarbon degradation | 121 | 2254 | 0.7537280966208090 | 950.0 | 55356.0 | 340.0 | 264.0 |
| 00625 Chloroalkane and chloroalkene degradation | 11 | 345 | 0.6468011531234290 | 75.0 | 56552.0 | 54.0 | 29.0 |
| 00626 Naphthalene degradation | 69 | 1438 | 0.6357825704670690 | 404.0 | 55875.0 | 217.0 | 238.0 |
| 00627 Aminobenzoate degradation | 106 | 2056 | 0.6457810046377090 | 721.0 | 55168.0 | 357.0 | 411.0 |
| 00630 Glyoxylate and dicarboxylate metabolism | 78 | 1961 | 0.6352256427893340 | 553.0 | 55918.0 | 390.0 | 237.0 |
| 00633 Nitrotoluene degradation | 33 | 658 | 0.8075168569911570 | 244.0 | 56672.0 | 20.0 | 101.0 |
| 00640 Propanoate metabolism | 41 | 1216 | 0.5275516914956930 | 253.0 | 56251.0 | 284.0 | 169.0 |
| 00642 Ethylbenzene degradation | 20 | 488 | 0.4856692068933750 | 70.0 | 56518.0 | 53.0 | 98.0 |
| 00643 Styrene degradation | 27 | 628 | 0.5442834314614210 | 143.0 | 56398.0 | 90.0 | 151.0 |
| 00650 Butanoate metabolism | 43 | 1339 | 0.6337107369408950 | 278.0 | 56370.0 | 185.0 | 134.0 |
| 00660 C5-Branched dibasic acid metabolism | 40 | 872 | 0.5609187135182550 | 237.0 | 56121.0 | 225.0 | 145.0 |
| 00670 One carbon pool by folate | 42 | 1048 | 0.5127876754159670 | 242.0 | 56678.0 | 305.0 | 159.0 |
| 00680 Methane metabolism | 102 | 2534 | 0.7533470795904140 | 856.0 | 55473.0 | 424.0 | 140.0 |
| 00710 Carbon fixation in photosynthetic organisms | 30 | 571 | 0.6911517217454610 | 239.0 | 56530.0 | 141.0 | 74.0 |
| 00720 Carbon fixation pathways in prokaryotes | 48 | 1706 | 0.7372234377314740 | 403.0 | 56001.0 | 193.0 | 95.0 |
| 00730 Thiamine metabolism | 34 | 598 | 0.6351616397377490 | 209.0 | 56495.0 | 94.0 | 146.0 |
| 00740 Riboflavin metabolism | 36 | 928 | 0.6430606851335270 | 239.0 | 56313.0 | 120.0 | 143.0 |
| 00750 Vitamin B6 metabolism | 42 | 664 | 0.68878962694997 | 301.0 | 56341.0 | 132.0 | 137.0 |
| 00760 Nicotinate and nicotinamide metabolism | 77 | 1312 | 0.7566014310904930 | 609.0 | 55501.0 | 212.0 | 173.0 |
| 00770 Pantothenate and CoA biosynthesis | 53 | 975 | 0.8125071653405430 | 477.0 | 55856.0 | 163.0 | 59.0 |
| 00780 Biotin metabolism | 29 | 704 | 0.6510390615787090 | 218.0 | 56140.0 | 173.0 | 67.0 |
| 00785 Lipoic acid metabolism | 41 | 1160 | 0.6104343767640310 | 333.0 | 56235.0 | 337.0 | 106.0 |
| 00790 Folate biosynthesis | 81 | 1941 | 0.7820022413017530 | 663.0 | 55408.0 | 244.0 | 123.0 |
| 00791 Atrazine degradation | 22 | 267 | 0.8879918741580400 | 225.0 | 56344.0 | 40.0 | 17.0 |
| 00830 Retinol metabolism | 33 | 1005 | 0.6565859869886410 | 275.0 | 56211.0 | 271.0 | 44.0 |
| 00860 Porphyrin metabolism | 161 | 7551 | 0.7786835720932580 | 1439.0 | 54827.0 | 577.0 | 225.0 |
| 00900 Terpenoid backbone biosynthesis | 58 | 1676 | 0.6983224177776770 | 470.0 | 55767.0 | 285.0 | 124.0 |
| 00901 Indole alkaloid biosynthesis | 96 | 2470 | 0.8787511753723070 | 887.0 | 55879.0 | 154.0 | 87.0 |
| 00902 Monoterpenoid biosynthesis | 70 | 1166 | 0.5985305822686890 | 412.0 | 55937.0 | 285.0 | 257.0 |
| 00903 Limonene degradation | 56 | 1156 | 0.8168080310251450 | 433.0 | 55751.0 | 97.0 | 95.0 |
| 00904 Diterpenoid biosynthesis | 135 | 3445 | 0.8595465627286350 | 1181.0 | 55543.0 | 237.0 | 140.0 |
| 00905 Brassinosteroid biosynthesis | 33 | 1088 | 0.9358212201321070 | 307.0 | 56710.0 | 28.0 | 14.0 |
| 00906 Carotenoid biosynthesis | 142 | 5553 | 0.8970744191367720 | 1348.0 | 55336.0 | 201.0 | 101.0 |
| 00907 Pinene, camphor and geraniol degradation | 65 | 2056 | 0.7458763981009230 | 497.0 | 56128.0 | 200.0 | 135.0 |
| 00908 Zeatin biosynthesis | 38 | 964 | 0.7921510901356950 | 312.0 | 56427.0 | 98.0 | 65.0 |
| 00909 Sesquiterpenoid and triterpenoid biosynthesis | 92 | 2033 | 0.6272931720065450 | 635.0 | 55392.0 | 417.0 | 319.0 |
| 00910 Nitrogen metabolism | 15 | 453 | 0.6766126048746800 | 96.0 | 56451.0 | 55.0 | 37.0 |
| 00920 Sulfur metabolism | 52 | 1345 | 0.6747486742538950 | 332.0 | 55763.0 | 162.0 | 154.0 |
| 00930 Caprolactam degradation | 35 | 827 | 0.5605678597768910 | 201.0 | 56555.0 | 186.0 | 128.0 |
| 00940 Phenylpropanoid biosynthesis | 74 | 1812 | 0.7649315369779430 | 618.0 | 55988.0 | 238.0 | 138.0 |
| 00941 Flavonoid biosynthesis | 92 | 2361 | 0.7736911928778960 | 761.0 | 55662.0 | 287.0 | 153.0 |
| 00942 Anthocyanin biosynthesis | 79 | 3666 | 0.8521201133396060 | 702.0 | 55893.0 | 155.0 | 86.0 |
| 00943 Isoflavonoid biosynthesis | 76 | 1902 | 0.8394004082119710 | 665.0 | 56336.0 | 149.0 | 102.0 |
| 00944 Flavone and flavonol biosynthesis | 66 | 2332 | 0.8267688659404890 | 637.0 | 56071.0 | 210.0 | 60.0 |
| 00945 Stilbenoid, diarylheptanoid and gingerol biosynthesis | 34 | 1108 | 0.75979790751362 | 263.0 | 56251.0 | 110.0 | 57.0 |
| 00946 Degradation of flavonoids | 47 | 1048 | 0.48029316462370500 | 239.0 | 56494.0 | 286.0 | 224.0 |
| 00950 Isoquinoline alkaloid biosynthesis | 144 | 3669 | 0.9095550637821420 | 1371.0 | 55811.0 | 148.0 | 117.0 |
| 00960 Tropane, piperidine and pyridine alkaloid biosynthesis | 91 | 1891 | 0.793513754160815 | 730.0 | 55647.0 | 213.0 | 160.0 |
| 00965 Betalain biosynthesis | 32 | 895 | 0.70684621101882 | 215.0 | 56478.0 | 87.0 | 90.0 |
| 00966 Glucosinolate biosynthesis | 89 | 1622 | 0.8882234493070090 | 798.0 | 55783.0 | 149.0 | 51.0 |
| 00980 Metabolism of xenobiotics by cytochrome P450 | 115 | 2346 | 0.8206921470458940 | 897.0 | 55710.0 | 180.0 | 203.0 |
| 00981 Insect hormone biosynthesis | 33 | 950 | 0.770038639507383 | 309.0 | 55970.0 | 144.0 | 45.0 |
| 00982 Drug metabolism - cytochrome P450 | 94 | 1809 | 0.9115830208550640 | 845.0 | 55678.0 | 84.0 | 77.0 |
| 00983 Drug metabolism - other enzymes | 74 | 1677 | 0.832535891433835 | 590.0 | 56134.0 | 106.0 | 128.0 |
| 00984 Steroid degradation | 29 | 811 | 0.780509722686123 | 245.0 | 56261.0 | 76.0 | 61.0 |
| 00996 Biosynthesis of various alkaloids | 89 | 2537 | 0.6769750135188250 | 632.0 | 55466.0 | 351.0 | 241.0 |
| 00997 Biosynthesis of various other secondary metabolites | 82 | 1720 | 0.620256681819166 | 581.0 | 55712.0 | 467.0 | 240.0 |
| 00998 Biosynthesis of various antibiotics | 126 | 2586 | 0.7469649931826780 | 960.0 | 55172.0 | 320.0 | 311.0 |
| 00999 Biosynthesis of various plant secondary metabolites | 174 | 5573 | 0.7524500635441920 | 1378.0 | 54640.0 | 512.0 | 360.0 |
| 01040 Biosynthesis of unsaturated fatty acids | 80 | 4509 | 0.7931406916340190 | 765.0 | 55455.0 | 406.0 | 23.0 |
| 01051 Biosynthesis of ansamycins | 48 | 1614 | 0.7428984261088510 | 389.0 | 56201.0 | 157.0 | 110.0 |
| 01052 Type I polyketide structures | 25 | 1280 | 0.42387899618925300 | 87.0 | 56704.0 | 84.0 | 157.0 |
| 01053 Biosynthesis of siderophore group nonribosomal peptides | 29 | 809 | 0.6271569045656970 | 183.0 | 56929.0 | 101.0 | 115.0 |
| 01054 Nonribosomal peptide structures | 14 | 1062 | 0.6934693151337760 | 72.0 | 56579.0 | 6.0 | 66.0 |
| 01055 Biosynthesis of vancomycin group antibiotics | 26 | 909 | 0.4828061733893560 | 129.0 | 56358.0 | 156.0 | 119.0 |
| 01056 Biosynthesis of type II polyketide backbone | 30 | 1034 | 0.8639344763724760 | 268.0 | 56292.0 | 48.0 | 36.0 |
| 01057 Biosynthesis of type II polyketide products | 160 | 5716 | 0.9271790666316450 | 1429.0 | 55208.0 | 119.0 | 99.0 |
| 01059 Biosynthesis of enediyne antibiotics | 73 | 2401 | 0.7364613478070240 | 546.0 | 55934.0 | 200.0 | 184.0 |
| **01060 Biosynthesis of plant secondary metabolites** | 134 | 2511 | 0.576574875334981 | 998.0 | 53880.0 | 1125.0 | 355.0 |
| **01061 Biosynthesis of phenylpropanoids** | 105 | 2123 | 0.5836969274582290 | 779.0 | 55420.0 | 822.0 | 299.0 |
| **01062 Biosynthesis of terpenoids and steroids** | 96 | 2135 | 0.5477297998792720 | 641.0 | 55026.0 | 727.0 | 328.0 |
| **01063 Biosynthesis of alkaloids derived from shikimate pathway** | 142 | 2962 | 0.7421287384147680 | 1150.0 | 54903.0 | 546.0 | 240.0 |
| **01064 Biosynthesis of alkaloids derived from ornithine, lysine and nicotinic acid** | 63 | 993 | 0.7425623581569070 | 499.0 | 55896.0 | 220.0 | 124.0 |
| **01065 Biosynthesis of alkaloids derived from histidine and purine** | 34 | 565 | 0.7307892738408580 | 280.0 | 56389.0 | 141.0 | 67.0 |
| **01066 Biosynthesis of alkaloids derived from terpenoid and polyketide** | 47 | 1161 | 0.5083709282611360 | 226.0 | 56450.0 | 201.0 | 230.0 |
| **01070 Biosynthesis of plant hormones** | 66 | 1454 | 0.46495064652427200 | 396.0 | 55407.0 | 724.0 | 231.0 |

